# Cold, dark, and hungry: Dynamic transcriptional regulation across eight months of brumation

**DOI:** 10.64898/2026.06.11.731677

**Authors:** David L. Hubert, Ehren J. Bentz, Robert T. Mason

## Abstract

Long-term winter dormancy in ectotherms (brumation) defines the annual cycle of many temperate-zone reptiles, yet the transcriptional regulation that supports survival across months of cold and aphagy remains poorly understood. We generated time-resolved transcriptomic profiles of liver and testis from male red-sided garter snakes (*Thamnophis sirtalis parietalis*) at five timepoints spanning the eight-month brumation cycle: pre-brumation, early, mid-, and late brumation, and post-arousal under continued aphagy. Time-course negative-binomial regression (maSigPro) followed by gene-set enrichment analysis identified 3,715 transcripts in liver and 5,828 in testis with significant temporal expression structure organized into five overarching temporal patterns: sustained downregulation, downregulation with post-arousal recovery, sustained upregulation, brumation-specific upregulation and cyclic modulation. Liver showed coordinated upregulation of fatty acid mobilization enzymes (ATGL, FOXO1, PPARα, CPT1A) and gluconeogenic regulators (CREBBP, PCK1) coincident with sustained low temperatures. Additionally, low temperature transcriptional activity was suggestive of a shift toward hepatic lipid mobilization and alanine-supported gluconeogenesis. Testis showed sustained suppression of meiosis, reproduction, and DNA-metabolism gene sets that did not fully recover at arousal consistent with this species’ dissociated reproductive pattern. Both tissues showed coordinated upregulation of stress-response pathways involving heat-shock proteins, HIF1α and a glutathione-based antioxidant defense. Interestingly, three vitellogenin transcripts and 17β-hydroxysteroid dehydrogenases associated with estradiol-favoring steroid metabolism were upregulated in male liver during late brumation, which is not expected during natural physiology in adult males. Together these data support a framework in which temperature- and starvation-associated transcriptional programs contribute to survival of one of the longest, coldest brumations documented in a squamate.

**Summary statement:** A time-resolved transcriptomic analysis of liver and testis spanning eight months of winter brumation in *Thamnophis sirtalis parietalis* reveals gene expression patterns consistent with a temperature-associated shift toward hepatic lipid mobilization, sustained reproductive suppression, and vitellogenin response in males.

## Introduction

Long-term dormancy in ectotherms has traditionally been framed as a problem of endurance, in which animals survive cold and starvation by doing as little as possible. To suggest that such prolonged dormancy might involve dynamic, coordinated, tissue-specific transcriptional activity would have seemed implausible a decade ago, yet the tools to test this idea are now practical. Seasonal dormancy is a widespread strategy employed by organisms across the tree of life to survive extremes of temperature and resource scarcity. Although these states are often collectively referred to as “hibernation”, the underlying physiology varies substantially among taxa and across environments and can be more usefully partitioned into estivation, diapause, hibernation, and brumation (Wilsterman et al., 2021). Here we use hibernation exclusively for endothermic dormancy and brumation for the long-term, low-temperature dormancy of ectotherms, in which environmental temperature directly governs body temperature and metabolic rate (Mayhew, 1965; Voituron et al., 2000; Storey, 2003).

Hibernation in mammals has been studied in detail and is characterized by long bouts of torpor, in which body temperature and metabolism can fall by ≥80–90% (Heldmaier et al., 2004). Importantly, hibernation is defined by punctuated brief periods of interbout arousal during which transcription, translation, drinking, waste elimination and parturition occur (French, 1985; Storey, 2003; Geiser, 2013; Ruf and Geiser, 2015). Mammalian hibernators typically upregulate transcripts that promote lipid utilization and that suppress carbohydrate use, transcription and the cell cycle (Andrews et al., 1998; Hittel and Storey, 2002; Carey et al., 2003; Andrews, 2019). Crucially, ectothermic brumators have no equivalent of the interbout arousal: body temperature tracks ambient temperature continuously, and any maintenance transcriptional activity must occur at temperatures that mammalian hibernators rarely encounter (Gregory, 1982; Geiser, 2013).

Despite this, transcriptomic studies of brumation are sparse. The first detailed survey of gene expression in a brumating ectotherm (the bearded dragon, *Pogona vitticeps*) compared a single late-brumation timepoint with two post-arousal timepoints and found >2,400 differentially expressed genes across brain, heart and muscle, with enrichment for chromatin remodeling, translation, ubiquitination, stress response, cell-cycle suppression, DNA damage repair and carbohydrate metabolism (Capraro et al., 2019). Subsequent work in the Chinese alligator (*Alligator sinensis*; Lin et al., 2020), the Chinese soft-shelled turtle (*Pelodiscus sinensis*; Tang et al., 2021) and miRNAs of bearded dragon (Capraro et al., 2020) has extended the picture, but each study examines only one or two brumation timepoints, leaving the temporal sequence of transcriptional events during brumation largely unexplored.

The red-sided garter snake (*Thamnophis sirtalis parietalis*) is a uniquely well-suited system in which to address this gap. This species follows an annual cycle divided into three discrete phases: winter brumation, spring mating, and summer feeding, with brumation spanning up to eight months at body temperatures of 2-4 °C in subterranean hibernacula (Aleksiuk and Stewart, 1971; Gregory, 1974; Lutterschmidt et al., 2006). Animals enter brumation aphagic and remain so through emergence and the subsequent month of intense scramble-mating before migrating ∼17 km to summer feeding grounds (O’Donnell et al., 2004). Unlike most squamates, several species of snake, including *T. s. parietalis* show no consistent measurable depletion of adipocyte lipid stores across brumation, and previous work has hypothesized that hepatic and muscle glycogen and protein, rather than fat, supply the bulk of energy during this period (Costanzo, 1985; Zani et al., 2012; Wilson, 2020). This discrepancy in metabolic substrate use remains unresolved, and transcript-level analysis offers a way to clarify which metabolic pathways are maintained, suppressed, or reactivated across the brumation cycle.

Here, we report a time-course transcriptomic study of liver and testis in adult male *T. s. parietalis* at five points spanning brumation: pre-brumation, early brumation, mid-brumation, late brumation, and post-arousal. The liver was selected because of its role as the central organ of energy partitioning and was anticipated to show coordinated metabolic remodeling. Testis was selected because of the species’ dissociated reproductive pattern, in which spermatogenesis is completed in late summer and the resulting sperm are stored over winter for use in the energetically demanding spring mating period (Crews et al., 1984; Krohmer et al., 1987; Clesson et al., 2002). Three questions motivated this study: (i) is the long brumation of *T. s. parietalis* characterized by simple global transcriptional suppression, or does it exhibit temporally structured programs?; (ii) which metabolic substrates are mobilized in the liver during brumation given the apparent absence of adipocyte depletion?; and (iii) is transcriptional activity in testis consistent with a passive sperm-storage role, or does testis tissue activate maintenance or stress pathways at temperatures near 0 °C?

We used time-course-aware negative-binomial regression (maSigPro; Conesa and Nueda, 2022) to identify clusters of co-regulated transcripts and functionally characterized clusters using gene-set enrichment analyses. We found that while overall metabolism was broadly depressed during brumation, both tissues mounted dynamic, tissue-specific transcriptional responses to low temperatures and prolonged aphagy. In liver, temporal expression patterns are consistent with a temperature-associated shift in metabolic substrate use, including upregulation of hepatic lipid mobilization and alanine-supported gluconeogenesis at the coldest timepoints, alongside a coordinated stress response in which antioxidant defense shifts from catalase- to glutathione-based pathways. In testis, sustained suppression of meiotic and reproductive gene sets across brumation is consistent with the species’ dissociated reproductive pattern, while brumation-specific upregulation of translation and RNA-processing machinery suggests active maintenance of stored sperm viability at near-freezing temperatures. Together, these findings reframe brumation in *T. s. parietalis* from a period of passive endurance to one of coordinated, tissue-specific molecular management.

## Materials and methods

### Animal collection and husbandry

All experimental procedures were approved by the Oregon State University Institutional Animal Care and Use Committee (ACUP 4818) and conducted under Manitoba Wildlife Scientific Permit WB20333. Adult male *T. s. parietalis* (n = 80) were collected by hand at communal hibernacula in the Interlake region of Manitoba, Canada, during the spring mating aggregation of 2018 and transported to Oregon State University in a thermally regulated vehicle (∼22 °C, ∼48 h transit). Adult males were selected based on estimated age, inferred from body size (SVL ∼49 cm, mass ∼37 g), apparent overall health, and behavioral observations confirming active courtship. The extremely large population sizes at the dens allowed selective collection of healthy, reproductively active males even at high sample sizes. Snakes were housed individually in 45 L aquaria with paper substrate, water *ad libitum*, a hide and a basking lamp, in microprocessor-controlled environmental chambers. From June through late September, photophase / scotophase temperatures were 22 / 16 °C and animals were fed a mixture of earthworms and salmon fry to satiation weekly. Food was withheld during the data-collection window, consistent with the species’ aphagic mating and brumation seasons (Crews et al., 1987; O’Donnell et al., 2004).

### Brumation conditions and tissue collection

Brumation was simulated based on field temperature data from natural hibernacula (Lutterschmidt et al., 2006). Beginning in the last week of September, photoperiod and temperature were ramped down until complete darkness and a sustained chamber temperature of 4 °C were reached in the first week of December; this regime was held until early April, when temperatures and photoperiod were ramped up to a photophase / scotophase of 15 / 11 °C (Fig. 1, Table S1). Animals remained unfed throughout the entire brumation and post-arousal sampling period, such that post-arousal individuals experienced restored active-season temperatures with continued aphagy. Individuals selected for tissue collection were determined by randomizing animal ID numbers, with eight animals sampled from the larger cohort at each of five timepoints to provide sufficient replication for detecting major transcriptional responses across the brumation time course (Schurch et al., 2016). The initial cohort size (n = 80) was chosen so that each timepoint, including the final post-arousal sample, could be randomly selected from a larger pool of available animals rather than being constrained to the remaining individuals. Sampling regime included 5 timepoints across the brumation cycle described in wild populations: pre-brumation (24 September), early brumation (20 November), mid-brumation (16 January), late brumation (17 April) and post-arousal (29 April; Fig. 1). Animals were euthanized by lethal subcutaneous Brevital® overdose (methohexital sodium, 0.005 ml g ¹, 1% solution), confirmed by absence of snout-tap reflex, and decapitated (Hubert et al. 2026). Liver and testis were dissected from each animal and immersed in RNAlater® within 5 minutes of removal from brumation conditions. Samples were held at 4 °C for 24 h, then stored at -20 °C until RNA extraction.

**Figure 1.**
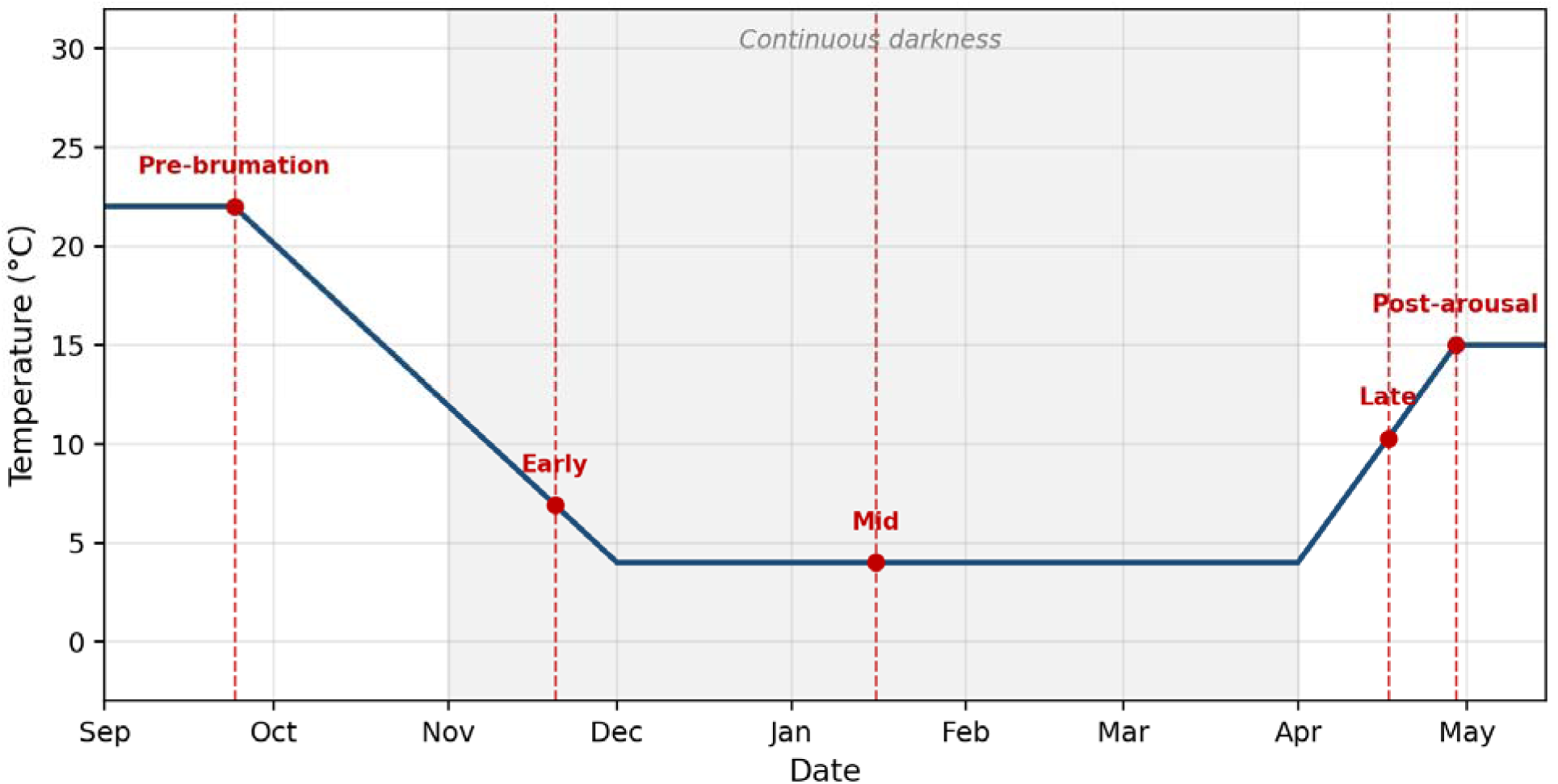
Experimental design. Brumation chamber temperature profile and tissue-collection schedule used in this study. Photophase / scotophase temperatures were ramped from active-season (22 / 16 °C) values down to a constant 4 °C in continuous darkness from December through early April, then ramped up to 15 / 11 °C; this profile was modelled on field measurements of body temperature in natural hibernacula (Lutterschmidt et al., 2006). Eight male *T. s. parietalis* were sampled at each of five timepoints (red dashed lines): pre-brumation, early brumation, mid-brumation, late brumation and post-arousal. Animals remained unfed throughout the experimental period. Liver and testis tissues are reported here.

### RNA extraction, library preparation, and sequencing

Total RNA was extracted from approximately 10 mg of liver or testis tissue using the E.Z.N.A.® HP Total RNA kit (Omega Bio-Tek). RNA was visualized using agarose gel electrophoresis with SYBR™ Safe staining, quantified using a Qubit™ 3 with RNA Broad Range kit, and diluted to a standardized concentration of 100 ng/ul. Libraries (n = 80) were constructed from 500 ng of total RNA using the Lexogen® QuantSeq 3′ mRNA-Seq library prep kit with sample-specific 6-nt i7 barcodes (Lexogen i7 Index Set 7001–7096). Optimal amplification cycle number (16) was determined by qPCR with the Lexogen PCR Add-on kit. Pooled libraries containing equal concentrations per sample were sequenced as 100-bp single-end reads on a single P2 flow cell of an Illumina® NextSeq™ 2000 at the Center for Quantitative Life Sciences, Oregon State University.

### Annotation with GOAnnotate

Transcripts from the *Thamnophis sirtalis parietalis* transcriptome assembly (NCBI BioProject PRJNA1357357) were functionally annotated using GOAnnotate (https://github.com/ehrenbentz/GOAnnotate). Translated transcript sequences were searched against the UniProt SwissProt and TrEMBL databases (release 2026_01) using Diamond BLASTx v2.1.9 (Buchfink et al., 2021) with an e-value threshold of 1e-4 and a maximum of 50 hits per query per database. SwissProt and TrEMBL were searched separately to ensure representation from both curated and computationally annotated protein databases, and the results were concatenated prior to annotation. GOAnnotate clusters BLAST hits by protein name similarity using average-linkage agglomerative clustering with a similarity threshold of 0.5, selects the best-supported cluster for each transcript, and assigns a protein name and gene symbol based on bitscore-weighted scoring within the winning cluster. 95.9% of annotations were based on best BLASTx hits from the TrEMBL database, and 4.1% from the SwissProt database. Non-redundant Gene Ontology terms were collected from the top 5 hits in the winning cluster and supplemented with terms from matching SwissProt entries, then filtered to the most specific terms using the GO hierarchy (go.obo; 2026_03_25). The resulting annotations yielded 32,639 transcripts annotated with protein names, 30,952 of which were assigned gene symbols, and 30,361 were assigned GO terms.

### Read processing and mapping

Reads were trimmed with bbduk (BBTools): the first 12 bp were clipped, residual adapter and low-quality termini were removed, reads <50 bp were discarded, and terminal stretches of A/G ≥5 bp were trimmed (Bushnell, 2014). Filtered reads were mapped to a multi-tissue annotated *T. s. parietalis* reference transcriptome with --qv-offset 33 -Q --strata -o 3 -N 4 -K 10000 -L (Hubert et al., 2026) using gmapper (SHRiMP2 v2.2.3; Rumble et al., 2009; David et al., 2011). Alignments were filtered to a minimum of 57 matches with SAMFilterByGene.pl and counts compiled with CombineExpression.pl (Meyer, 2018).

### Time-series differential expression and clustering

Counts were imported into R 4.2 and filtered to retain transcripts with an average of ≥5 reads per sample. Time-series differential expression was evaluated with maSigPro v1.68.0 using time-course-aware negative-binomial regression on TMM-normalized counts (Conesa and Nueda, 2022; Tarazona et al., 2011). Timepoint was modeled as a continuous variable using a fourth-degree polynomial, corresponding to the maximum polynomial degree supported by the five sampled timepoints. This approach allowed maSigPro to capture nonlinear expression trajectories across the brumation cycle, including monotonic changes, mid-brumation peaks, post-arousal recovery, and cyclic patterns. Transcripts with adjusted p < 0.05 in stepwise regression were grouped into 18 hierarchical clusters (k = 18), per analysis (combined liver + testis, liver-only, testis-only). The number of clusters was evaluated independently for each analysis (combined, liver-only and testis-only) using mean silhouette width (R package cluster; Rousseeuw, 1987) across k = 8–28 (Fig. S1). Silhouette scores declined after an initial inflection at k = 10, 11 and 12 for the testis, combined and liver analyses, respectively, followed by a stable plateau through k = 18 before declining further. We selected k = 18 as the minimum partition that (i) fell consistently within the stable plateau across all three analyses, (ii) yielded smaller mean cluster sizes (compared to smaller values of k) of 324, 206 and 627 transcripts for the testis, liver and combined analyses, respectively, and (iii) resolved biologically distinct expression sub-programs that collapsed into broader, less interpretable clusters at lower values of k. Multiple-test correction used the Benjamini–Hochberg procedure throughout.

### Gene-set enrichment

Gene Ontology enrichment analysis was performed using DEGOE (https://github.com/ehrenbentz/DEGOE). For each co-expression cluster identified by maSigPro, genes were classified as cluster members (1) or background consisting of all transcripts assigned to any cluster (0). Enrichment of GO terms among cluster members was tested using one-sided Fisher’s exact tests for over-representation within each cluster. GO annotations for each transcript were obtained from the gene2go.tsv generated by the GOAnnotate pipeline. The Gene Ontology hierarchy definitions file (go.obo; 2026_03_25) was used to propagate annotations from each directly annotated term to all ancestor terms via both is_a and part_of relationships.

GO categories were filtered to exclude overly broad terms: Biological Process (BP) categories below hierarchy level 3 and Molecular Function (MF) categories below level 2 were removed, and no level filter applied to Cellular Component (CC). Categories containing fewer than 5 genes or more than 10% of the total annotated gene set were excluded from analyses. Enrichment was tested separately for each cluster within each GO namespace (BP, MF, CC) and for each tissue (testis, liver, and combined). P-values were corrected for multiple testing using Benjamini-Hochberg FDR and GO terms with an adjusted p-value below 0.05 were considered significant.

## Results

### Sequencing, mapping and overall expression

A single P2 NextSeq 2000 lane yielded 540.7 × 10 reads, of which 528 × 10 were demultiplexed to one of the 80 libraries (mean 6.6 ± 0.45 × 10 reads sample ¹; mean Phred score 32.9). After filtering (mean 6.0 ± 0.42 × 10 reads sample ¹) and mapping, an average of 4.1 ± 0.28 × 10 reads sample ¹ mapped to the reference transcriptome (Table S2).

### Time-course clustering

Time-course-aware regression with maSigPro identified 11,286 transcripts with significant temporal structure in the combined analysis, 3,715 in liver and 5,828 in testis after stepwise model selection (padj < 0.05). Hierarchical clustering produced 18 clusters per analysis (Fig. 2-3). The combined-tissue analysis cleanly separated tissue-specific clusters from clusters with concordant changes in both tissues, identifying genes that respond uniformly to brumation as well as genes whose response is restricted to one tissue (Fig. S2).

**Figure 2.**
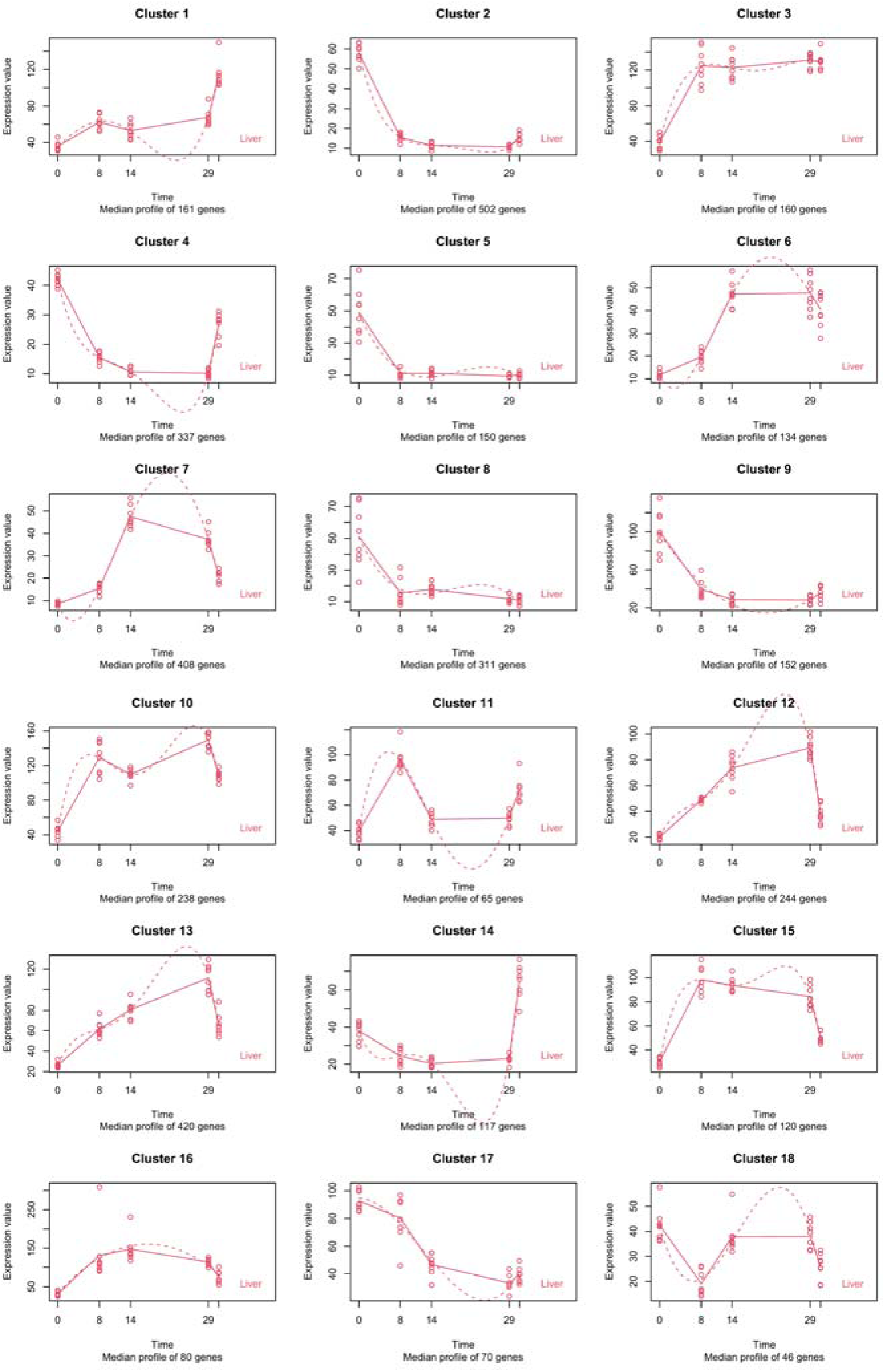
Liver temporal expression clusters across brumation. Hierarchical clustering of transcripts with significant temporal structure of 18 clusters in the liver analysis. Median-profile plots are shown for each cluster. The x-axis is time in weeks since the start of brumation; the post-arousal timepoint (31 weeks) is shown to the right of each panel. Points represent normalized expression values for individual biological replicates, with each biological replicate corresponding to one adult male snake; n = 8 animals per timepoint. The experiment was performed once as a time-course RNA-seq experiment using independently sampled animals at each timepoint. Solid lines are observed median expression profiles; dotted lines are fitted regression models. Clusters were identified using time-course-aware negative-binomial regression in maSigPro on TMM-normalized counts, with Benjamini–Hochberg correction for multiple testing; transcripts shown were significant at adjusted P < 0.05.

**Figure 3.**
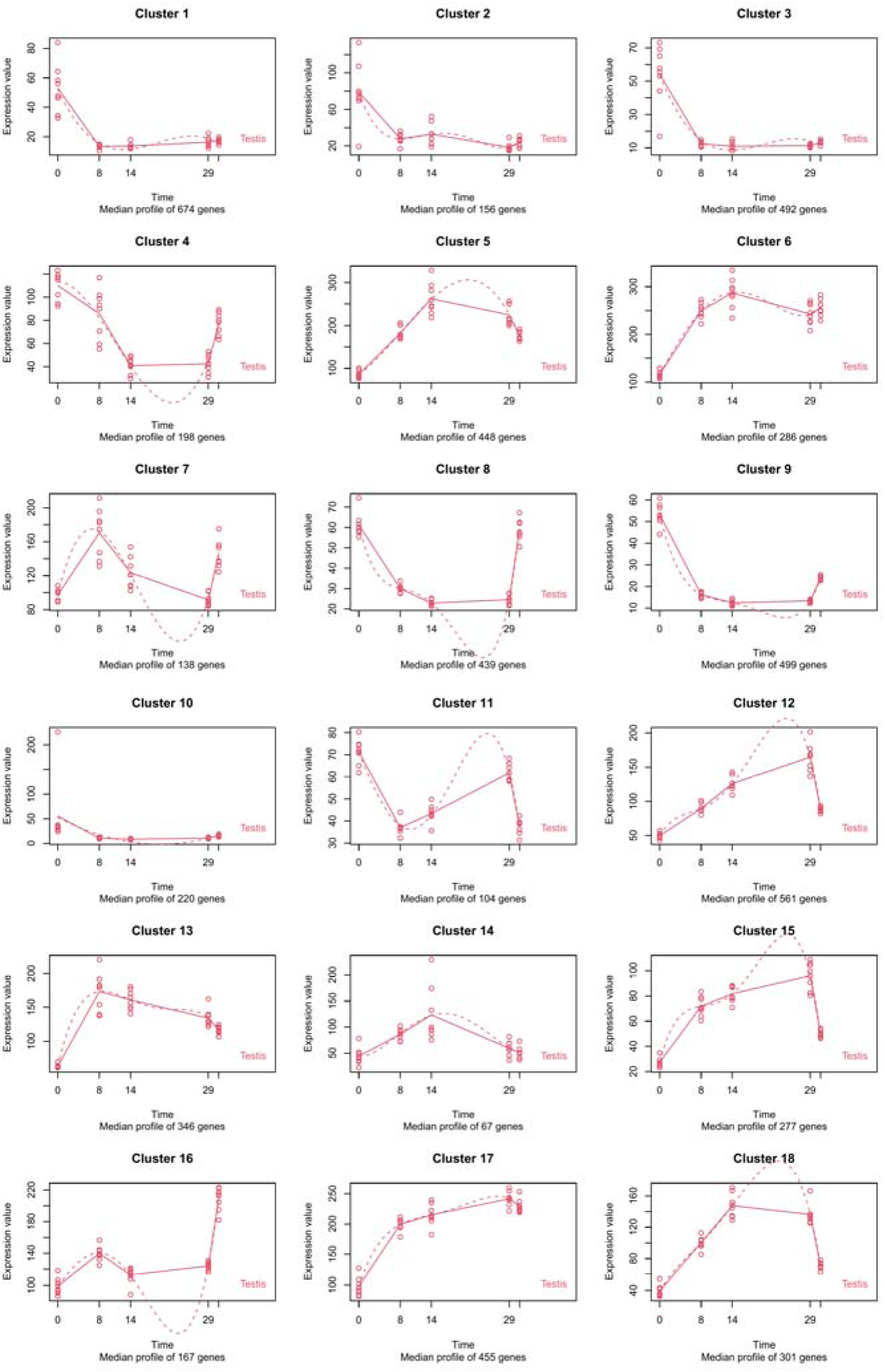
Testis temporal expression clusters across brumation. Hierarchical clustering of transcripts with significant temporal structure of 18 clusters in the testis analysis. Median-profile plots are shown for each cluster. The x-axis is time in weeks since the start of brumation; the post-arousal timepoint (31 weeks) is shown to the right of each panel. Points represent normalized expression values for individual biological replicates, with each biological replicate corresponding to one adult male snake; n = 8 animals per timepoint. The experiment was performed once as a time-course RNA-seq experiment using independently sampled animals at each timepoint. Solid lines are observed median expression profiles; dotted lines are fitted regression models. Clusters were identified using time-course-aware negative-binomial regression in maSigPro on TMM-normalized counts, with Benjamini–Hochberg correction for multiple testing; transcripts shown were significant at adjusted P < 0.05.

Across all three maSigPro analyses, the 18 clusters were broadly assignable to five overarching expression archetypes (Fig. 4) based on the direction and recovery pattern of their median expression profiles: (i) sustained downregulation during brumation that did not recover post-arousal, (ii) downregulation with post-arousal recovery, (iii) sustained upregulation that persisted post-arousal, (iv) upregulation during brumation that decayed at arousal, and (v) cyclic modulation with one or more reversals during the cycle.

**Figure 4.**
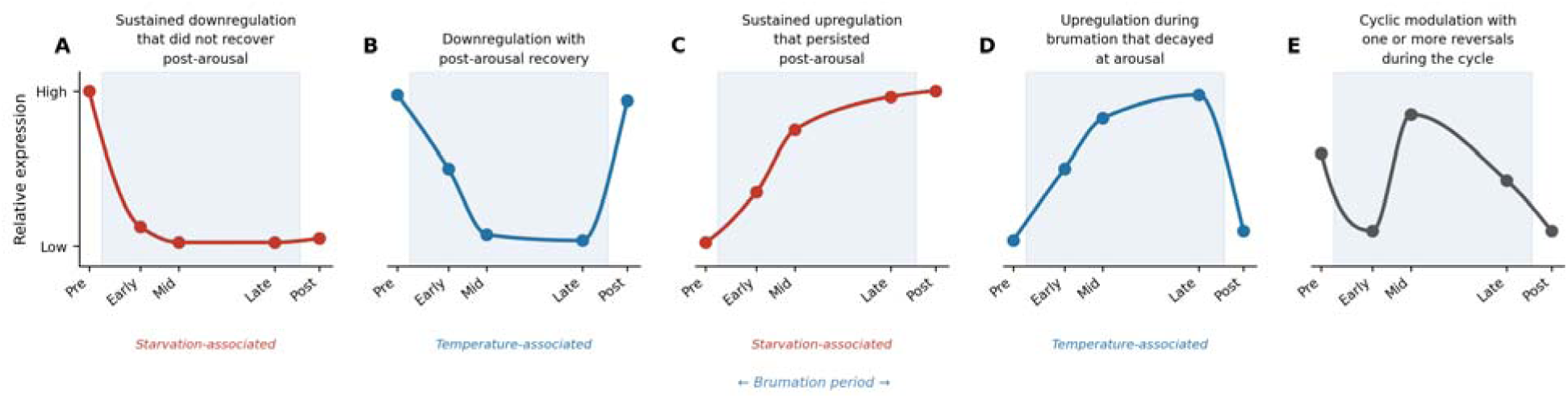
Five generalized expression archetypes that emerge from the time-course analysis. Schematic representations of the major patterns observed across clusters in both tissues: (A) sustained downregulation, (B) downregulation with post-arousal recovery, (C) sustained upregulation, (D) brumation-specific upregulation, (E) cyclic modulation. Patterns (A) and (C) track prolonged starvation; (B) and (D) track temperature.

### Functional Enrichment

Liver showed gene-set enrichment in 10 of 18 clusters (Table 1, Table S3). Three sustained-downregulation clusters were enriched: cluster 2 for lipid and fatty acid catabolic processes, organic acid catabolism, and oxidoreductase activity; cluster 8 for DNA metabolic process and base-excision repair; and cluster 9 for oxidative phosphorylation, the mitochondrial electron transport chain, ATP synthesis coupled electron transport, and respiratory complex I, spanning all three GO ontology domains (75 significant gene sets). Median expression of cluster 9 dropped >70% during brumation and did not recover. A single sustained-upregulation cluster (cluster 1) was enriched for ER unfolded protein response, response to ER stress, proteasome complex and regulatory particle, regulation of proteasomal protein catabolic process, and response to oxidative stress and type I interferon, indicating maintained protein quality control and innate immune capacity across brumation. Cluster 4, showing downregulation with post-arousal recovery, was enriched for cytochrome complex assembly, respiratory chain complex III assembly, and mitochondrial translation. Among brumation-specific upregulation clusters, cluster 7 was enriched for immune response across 48 significant gene sets, including lymphocyte mediated immunity, complement activation, humoral immune response, and inflammatory response, as well as chromatin organization and remodeling; cluster 13 was enriched for transcription cis-regulatory region binding and DNA-binding transcription factor activity. Three additional clusters reached significance with four or fewer enriched terms each (cluster 5: intermediate filament cytoskeleton; cluster 11: Golgi membrane; cluster 15: melanosome).

**Table 1.** Selected enriched gene-set themes per cluster, by tissue. Enrichment tested with DEGOE (Fisher’s exact test, FDR < 0.05). Bold labels denote individual clusters; unlabelled rows apply to all listed clusters. Full results in Tables S3–S5.

| Tissue | Cluster(s) | Pattern | Enriched themes |
| --- | --- | --- | --- |
| Liver | 2, 8, 9 | *Sustained downregulation* | **Cluster 2:** lipid and fatty acid catabolic process; organic acid catabolism; oxidoreductase activity; peroxisome. **Cluster 8:** DNA metabolic process; base-excision repair; DNA biosynthesis; monoamine transport. **Cluster 9:** oxidative phosphorylation; mitochondrial electron transport chain; ATP synthesis coupled electron transport; respiratory complex I; mitochondrial envelope and membranes |
| Liver | 1 | *Sustained upregulation* | Response to oxidative stress; ER unfolded protein response; response to ER stress; proteasome complex; regulation of proteasomal protein catabolic process; response to type I interferon; regulation of AMPA receptor activity |
| Liver | 4 | *Brumation-specific downregulation* | Cytochrome complex assembly; respiratory chain complex III assembly; mitochondrial translation; mitochondrial matrix; mitochondrial respiratory chain complex III assembly |
| Liver | 7, 13 | *Brumation-specific upregulation* | **Cluster 7:** immune response; lymphocyte mediated immunity; complement activation; humoral immune response; inflammatory response; chromatin organization and remodeling; regulation of MAPK cascade; neurogenesis. **Cluster 13:** transcription cis-regulatory region binding; DNA-binding transcription factor activity; RNA polymerase II regulatory region binding; sequence-specific DNA binding; transcription factor binding |
| Liver | 5, 11, 15 | *Various* | **Cluster 5 (sustained-down):** intermediate filament cytoskeleton; postsynapse [minor, 4 terms]. **Cluster 11 (cyclic):** Golgi membrane [minor, 1 term]. **Cluster 15 (brumation-specific upregulation):** melanosome; pigment granule [minor, 2 terms] |
| Testis | 1, 2, 3, 10 | *Sustained downregulation* | **Cluster 1:** DNA metabolic process; base-excision repair; RNA-directed DNA polymerase activity; retrotransposition; sodium ion homeostasis. **Clusters 2, 3:** meiotic cell cycle; sperm axoneme assembly and flagellum motility; gamete generation; spermatogenesis; chromosome segregation; cilium organization; DNA recombination and repair. **Cluster 10:** leukocyte mediated immunity; B cell activation and proliferation; adaptive immune response; defense response; inflammatory response; leukocyte differentiation [179 enriched terms, largest cluster] |
| Testis | 6, 17 | *Sustained upregulation* | **Cluster 6:** ubiquitin-dependent protein catabolic process; proteasome-mediated ubiquitin-dependent protein catabolic process; regulation of catabolic process. **Cluster 17:** GTPase activity; G protein activity; GTP and guanyl nucleotide binding |
| Testis | 4, 8, 9 | *Brumation-specific downregulation* | **Cluster 4:** ribosomal subunit export from nucleus; ribosome localization; nuclear export; negative regulation of transcription and RNA metabolic process; DNA integration. **Cluster 8:** rRNA processing; rRNA metabolic process; ribosome biogenesis. **Cluster 9:** tissue homeostasis; cell projection organization; neuron differentiation; anatomical structure homeostasis |
| Testis | 5, 12, 13 | *Brumation-specific upregulation* | **Cluster 5:** cytoplasmic translation; ribosome biogenesis and assembly; ribonucleoprotein complex biogenesis; muscle contraction; oxidative phosphorylation [81 enriched terms]. **Cluster 12:** organellar ribosome; mitochondrial ribosome; mitochondrial matrix; GTPase activity. **Cluster 13:** mRNA processing; alternative mRNA splicing via spliceosome; mRNA metabolic process; positive regulation of cell differentiation; miRNA processing |
| Testis | 7 | *Cyclic modulation* | Nuclear retinoic acid receptor binding; ATPase binding [minor, 2 terms] |

Testis showed gene-set enrichment in 13 of 18 clusters, with cluster 10 carrying the highest number of significantly enriched gene sets (179 terms; Table 1, Table S4). Sustained-downregulation clusters (clusters 1, 2, 3, 10) were enriched for meiosis, reproduction, DNA metabolism, and immune response, and did not recover post-arousal, consistent with the regressed reproductive state of testis during the spring mating period (Crews *et al*., 1984; Krohmer *et al*., 1987). Clusters 2 and 3 carried the strongest reproductive signal, enriched for meiotic cell cycle, spermatogenesis, sperm axoneme assembly, gamete generation, and chromosome segregation. Cluster 1 was enriched for DNA metabolic process, base-excision repair, and RNA-directed DNA polymerase activity. Cluster 10 was enriched for adaptive and innate immune response, leukocyte and B cell activation, leukocyte differentiation, and defense response. Two sustained-upregulation clusters were also enriched: cluster 6 for ubiquitin-dependent protein catabolic process and proteasome-mediated ubiquitin-dependent protein catabolic process, and cluster 17 for GTPase activity and G protein activity. A second set of clusters (4, 8, 9) showed downregulation followed by partial post-arousal recovery; cluster 4 was enriched for ribosomal subunit export from nucleus, nuclear export, and negative regulation of transcription; cluster 8 for rRNA processing and ribosome biogenesis; and cluster 9 for tissue homeostasis and cell projection organization. Brumation-specific upregulation clusters (5, 12, 13) were enriched for translation, ribosome biology, and RNA processing: cluster 5 for cytoplasmic translation, ribosome biogenesis, and ribonucleoprotein complex biogenesis (81 significant gene sets; ∼250% peak expression relative to pre-brumation); cluster 12 for organellar ribosome, mitochondrial ribosome, and GTPase activity; and cluster 13 for mRNA processing, alternative mRNA splicing via spliceosome, and mRNA metabolic process.

The combined liver + testis analysis is presented as a broad comparison of shared versus tissue-specific temporal structure, and showed gene-set enrichment in 17 of 18 clusters (Table S5). Notably, combined-analysis cluster 1, the largest downregulated cluster, captures broad suppression of core metabolic processes and provides a reference background for interpreting more selective transcript-level changes during brumation. Median transcript abundance in this cluster fell approximately 70% from pre-brumation to mid-brumation, tracking the decline in chamber temperature, and partially recovered at post-arousal (Fig. S2). Gene-set enrichment of cluster 1 spanned virtually all branches of intermediary metabolism: oxidative phosphorylation, the mitochondrial electron transport chain and respiratory chain complexes I and IV, ATP biosynthesis, cellular respiration, fatty acid beta-oxidation, amino acid catabolism (including branched-chain and aromatic amino acid degradation), and carbohydrate catabolism (Table S5). The associated cellular component terms were dominated by mitochondrial inner membrane and peroxisome, consistent with suppression of both mitochondrial and peroxisomal oxidative capacity. This widespread transcriptional downregulation is consistent with a thermally-driven reduction in enzymatic demand at low body temperatures, and the cluster median is used throughout as a reference baseline against which the magnitude of change in specific transcripts of interest is compared.

### Lipid metabolism in liver suggests a coordinated cold-associated mobilization

Transcripts associated with hepatic triglyceride mobilization showed strong, temperature-tracking upregulation, despite broader downregulation of general metabolic pathways (Fig. 5A). Adipose triglyceride lipase (ATGL), the rate-limiting enzyme for hydrolysis of long-chain triglycerides stored in lipid droplets (Zimmermann et al., 2004; Smirnova et al., 2006), peaked at mid-brumation when chamber temperatures were lowest. Forkhead box O1 (FOXO1), a transcriptional activator of ATGL-dependent lipolysis (Chakrabarti and Kandror, 2009; Zhang et al., 2016), and peroxisome-proliferator-activated receptor α (PPARα), a master regulator of fatty acid oxidation and gluconeogenesis (Tyagi et al., 2011; Chiazza and Collino, 2016), followed the same pattern. Carnitine palmitoyltransferase 1A (CPT1A), required to import long-chain acyl-CoAs into the mitochondrion for β-oxidation (IJlst et al., 1998), showed an early-brumation expression spike followed by sustained moderate elevation, and CPT2, similarly required for β-oxidation (Gobin et al., 2003) was elevated throughout brumation.

**Figure 5.**
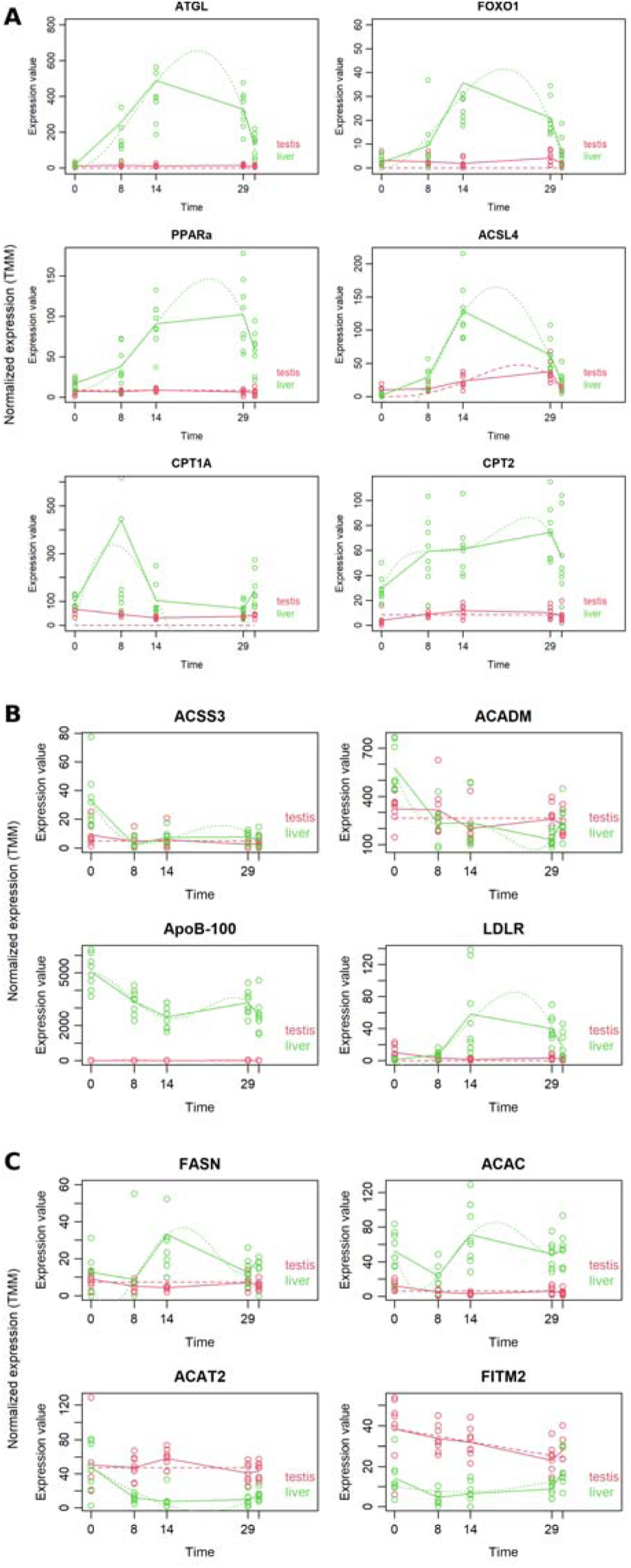
Coordinated hepatic lipid program during brumation. Liver (green) and testis (red) expression patterns are shown for transcripts associated with (A) lipid-droplet mobilization and long-chain fatty acid β-oxidation: ATGL, FOXO1, PPARα, ACSL4, CPT1A and CPT2; (B) substrate switching and lipoprotein transport: ACSS3, ACADM, ApoB-100 and LDLR; and (C) fatty acid biosynthesis and storage: FASN, ACAC, ACAT2 and FITM2. Points represent normalized expression values for individual biological replicates, with each biological replicate corresponding to one adult male snake; n = 8 animals per timepoint for each tissue. The experiment was performed once as a time-course RNA-seq experiment using independently sampled animals at each timepoint. Solid lines show observed mean expression, and dotted lines show fitted regression models. The x-axis is time in weeks from brumation onset; the post-arousal timepoint occurred at week 31 and is shown as the rightmost set of symbols in each panel. Transcripts were analyzed using time-course-aware negative-binomial regression in maSigPro on TMM-normalized counts with Benjamini–Hochberg correction for multiple testing; transcripts shown were significant at adjusted P < 0.05 in at least one analysis.

A complementary set of genes implicated a substrate switch within the fatty acid pool (Fig. 5A, B). Acyl-CoA synthetase long-chain family member 4 (ACSL4), which preferentially activates arachidonic acid released from triglycerides during ATGL-driven lipolysis (Ohkuni et al., 2013; Bermúdez et al., 2021), was strongly upregulated during mid-brumation. In contrast, ACSS3 (short-chain fatty acid activation; Yoshimura et al., 2017) and ACADM (medium-chain acyl-CoA dehydrogenase, the first step of medium-chain β-oxidation; Nandy et al., 1996) were downregulated through brumation.

Transport-related transcripts showed complementary patterns (Fig. 5B). Low-density lipoprotein receptor (LDLR) was strongly upregulated at mid-brumation, while apolipoprotein B-100 (ApoB-100), a structural component of hepatic VLDL output and the LDL-recognition motif (Olofsson et al., 1999), was reduced by only ∼40%, substantially less than the ∼70% reduction shown by general metabolic transcripts in the combined-cluster 1 (Fig. S2).

Surprisingly, lipid-biosynthetic enzymes also rose during brumation (Fig. 5C). Acetyl-CoA carboxylase 1 (ACAC) and fatty acid synthase (FASN) both peaked at mid-brumation. However, the lipid-droplet-storage enzymes ACAT2/SOAT2 (Ahmed et al., 2019; Pramfalk et al., 2022) and FITM2 (Kadereit et al., 2008; Becuwe et al., 2020) were downregulated.

### Carbohydrate metabolism in liver suggests alterations to glycogen metabolism, Cahill cycle activity, and gluconeogenesis during brumation

No liver cluster was enriched for upregulated carbohydrate-metabolism gene sets in our data. Glycogen branching enzyme (GBE1) and α-glucosidase (GAA), which respectively control glycogen synthesis branching and lysosomal glycogen catabolism, were both downregulated, but GAA decreased less steeply than the general metabolic background (Fig. 6A). Hexokinase 1 (HK1) was expressed at near-zero levels in liver throughout, while the hepatic glucose exporter GLUT2 showed a comparatively modest (∼45%) reduction (Fig. 6A).

**Figure 6.**
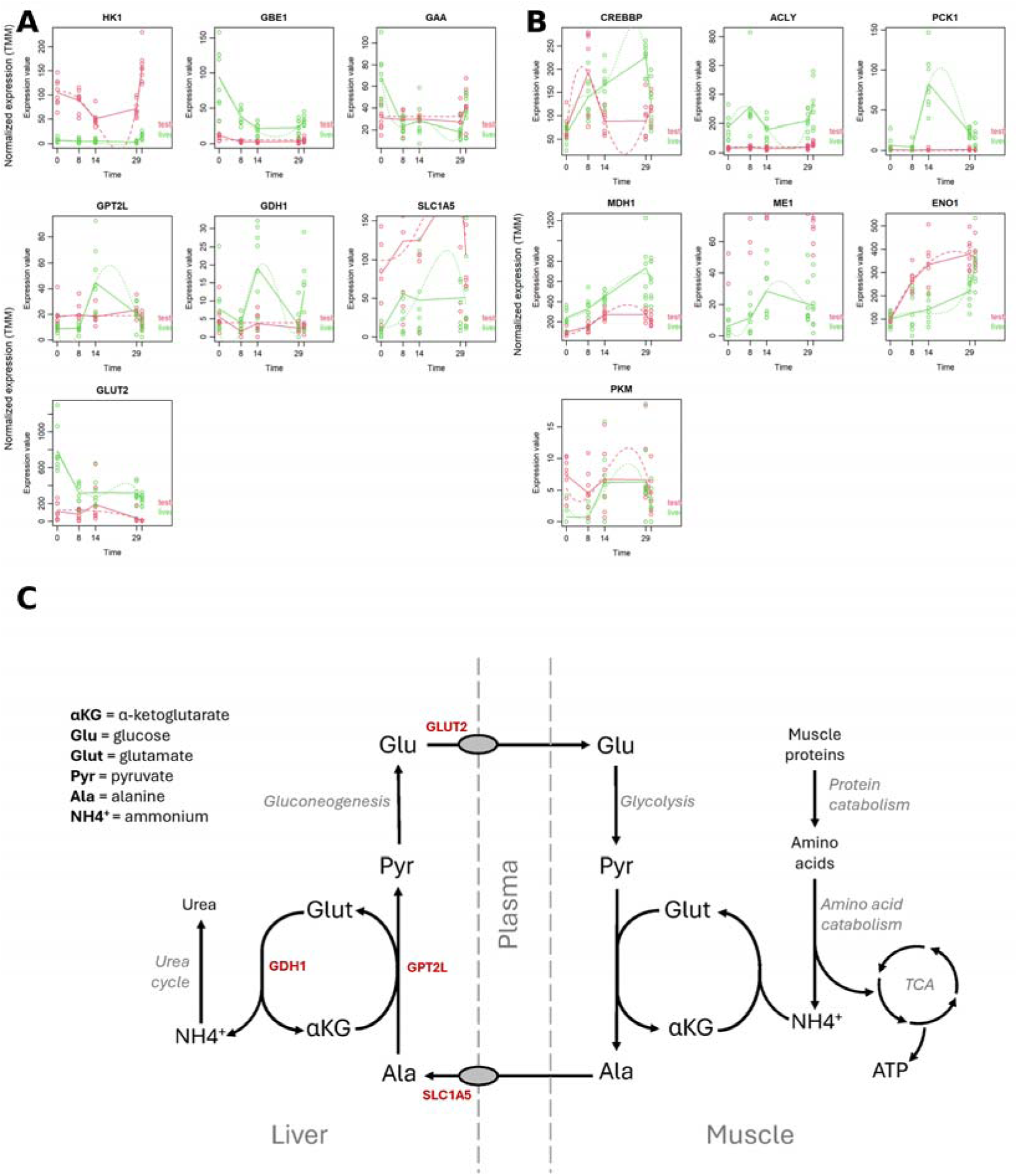
Hepatic glycogen handling, the Cahill cycle and gluconeogenesis. (A) Liver (green) and testis (red) expression patterns for glycogen and glucose-handling enzymes HK1, GBE1, GAA, alanine-cycle components GPT2L, GDH1, SLC1A5 and GLUT2. (B) Expression patterns for gluconeogenic regulators and enzymes: CREBBP, ACLY, PCK1, MDH1, ME1, ENO1 and PKM. Points represent TMM-normalized expression values for individual biological replicates, with each biological replicate corresponding to one adult male snake; n = 8 animals per timepoint for each tissue. The experiment was performed once as a time-course RNA-seq experiment using independently sampled animals at each timepoint. Solid lines show observed mean expression for each tissue/timepoint, and dotted lines show fitted regression models. The x-axis is time in weeks from brumation onset; the post-arousal timepoint occurred at week 31 and is shown as the rightmost set of symbols in each panel. Transcripts were analyzed using time-course-aware negative-binomial regression in maSigPro on TMM-normalized counts with Benjamini–Hochberg correction for multiple testing; transcripts shown were significant at adjusted P < 0.05 in at least one analysis. (C) Schematic of the glucose–alanine (Cahill) cycle linking muscle protein catabolism to hepatic gluconeogenesis. Genes labelled in red are those for which coordinated transcriptional change was observed.

In times of glucose scarcity, the Cahill (glucose–alanine) cycle uses alanine as a nitrogen carrier from skeletal muscle to liver with carbon recycled as glucose back to muscle (Felig, 1973; Sarabhai and Roden, 2019; Petersen et al., 2019; Fig. 6C). Hepatic transcripts glutamate dehydrogenase (GDH1) and alanine transaminase (GPT2L) showed strong upregulation peaking at mid-brumation alongside the lipid-mobilization panel (Fig. 6A). The neutral amino-acid transporter SLC1A5, which imports alanine and glutamine (Scalise et al., 2018), was sustained-upregulated through brumation and into post-arousal.

A similar mid-brumation peak was found in transcripts central to gluconeogenesis (Fig. 6B). FOXO1 (already implicated in lipolysis) and CREB-binding protein (CREBBP), both of which co-activate hepatic gluconeogenesis (Goodman and Smolik, 2000; Zhang et al., 2006), were strongly upregulated during brumation. Phosphoenolpyruvate carboxykinase 1 (PCK1), the irreversible step that funnels oxaloacetate into the gluconeogenic branch (Han et al., 2016), spiked sharply at mid-brumation and dropped at arousal. ATP-citrate lyase (ACLY), which provides cytosolic acetyl-CoA from mitochondrial citrate, malate dehydrogenase 1 (MDH1) and malic enzyme 1 (ME1) all rose during brumation, providing a plausible source of carbon for both gluconeogenesis and the lipid-biosynthetic surge described above. α-Enolase (ENO1) was sustained-upregulated across brumation (Fig. 6B).

### Stress responses are tissue-specific and integrated

A diverse stress-response program was observed in both tissues but with distinct emphases (Fig. 7). Heat shock protein (HSP) transcripts displayed differential tissue distribution: HSP90A, HSP105, HSC71 and to a lesser extent HSP70 were more strongly elevated in testis, whereas HSP40A and HSC90 were more strongly elevated in liver (Fig. 7A). Expression of most HSP transcripts dropped sharply at arousal, consistent with reduction of long-term proteomic stress as temperatures rose.

**Figure 7.**
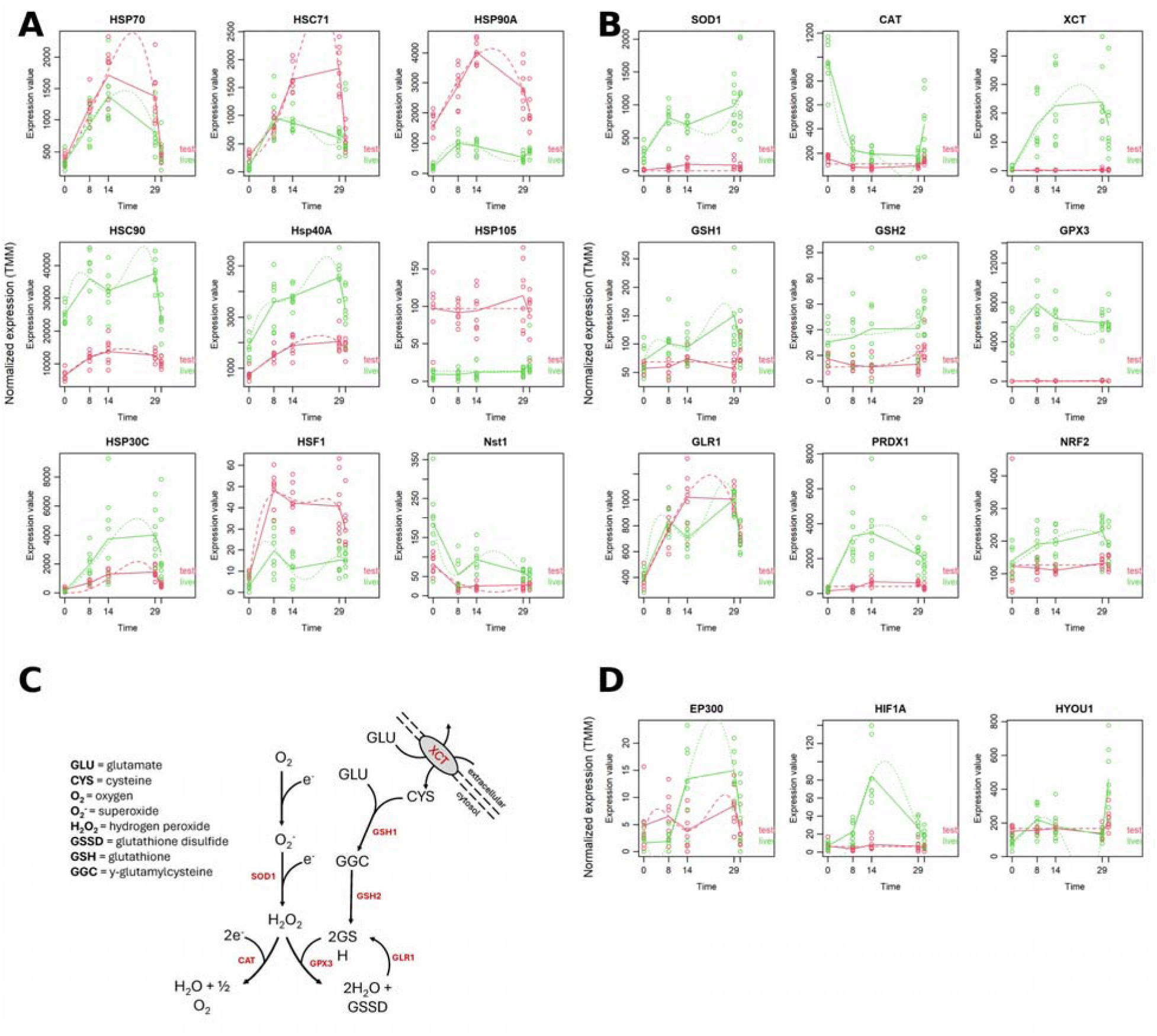
Integrated stress responses across brumation. (A) Liver (green) and testis (red) expression patterns for heat shock and chaperone-associated transcripts: HSP70, HSC71, HSP90A, HSC90, HSP40A, HSP105, HSP30C, HSF1 and Nst1. (B) Expression patterns for oxidative-stress-associated transcripts: SOD1, CAT, XCT, GSH1, GSH2, GPX3, GLR1, PRDX1 and NRF2. (C) Schematic of the oxidative-stress response showing the catalase and glutathione branches. Genes labelled in red are those for which coordinated transcriptional change was observed. (D) Expression patterns for hypoxia-associated transcripts: EP300, HIF1A and HYOU1. Points represent TMM-normalized expression values for individual biological replicates, with each biological replicate corresponding to one adult male snake; n = 8 animals per timepoint for each tissue. The experiment was performed once as a time-course RNA-seq experiment using independently sampled animals at each timepoint. Solid lines show observed mean expression for each tissue/timepoint, and dotted lines show fitted regression models. The x-axis is time in weeks from brumation onset; the post-arousal timepoint occurred at week 31 and is shown as the rightmost set of symbols in each panel. Transcripts were analyzed using time-course-aware negative-binomial regression in maSigPro on TMM-normalized counts with Benjamini–Hochberg correction for multiple testing; transcripts shown were significant at adjusted P < 0.05 in at least one analysis.

Transcripts associated with hypoxia response, including the transcriptional co-activator EP300, hypoxia-inducible factor 1α (HIF1A) and the ER-resident hypoxia-upregulated chaperone HYOU1, were upregulated during brumation in liver (Fig. 7D). EP300 and HIF1A peaked at mid-brumation and dropped at arousal (Wang et al., 1995; Pugh and Ratcliffe, 2003; Tamukong et al., 2022), while HYOU1 was sustained-upregulated and showed a further increase post-arousal (Ozawa et al., 1999).

The oxidative-stress response showed a striking switch between catalase- and glutathione-based pathways (Fig. 7B, C). Cytosolic superoxide dismutase (SOD1) was sustained-upregulated through brumation and peaked post-arousal. The glutathione synthase isoforms GSH1 and GSH2, glutathione peroxidase 3 (GPX3), glutathione reductase 1 (GLR1), peroxiredoxin 1 (PRDX1) and the cystine/glutamate antiporter XCT were upregulated during brumation, while catalase (CAT), which had high pre-brumation expression, was strongly downregulated and only partially recovered post-arousal. NRF2 was upregulated during brumation, indicating a broad transcriptional engagement of the antioxidant response (Ma, 2013).

### Evidence of estrogen modulation in male liver

Three vitellogenin (VTG) transcripts, vitellogenin-A2, vitellogenin-2 and a vitellogenin fragment, were strongly and concordantly upregulated in male liver beginning during late brumation and remained elevated post-arousal (Fig. 8A). The concordant upregulation of multiple VTG transcripts argues against annotation artifact and supports a genuine biological signal.

**Figure 8.**
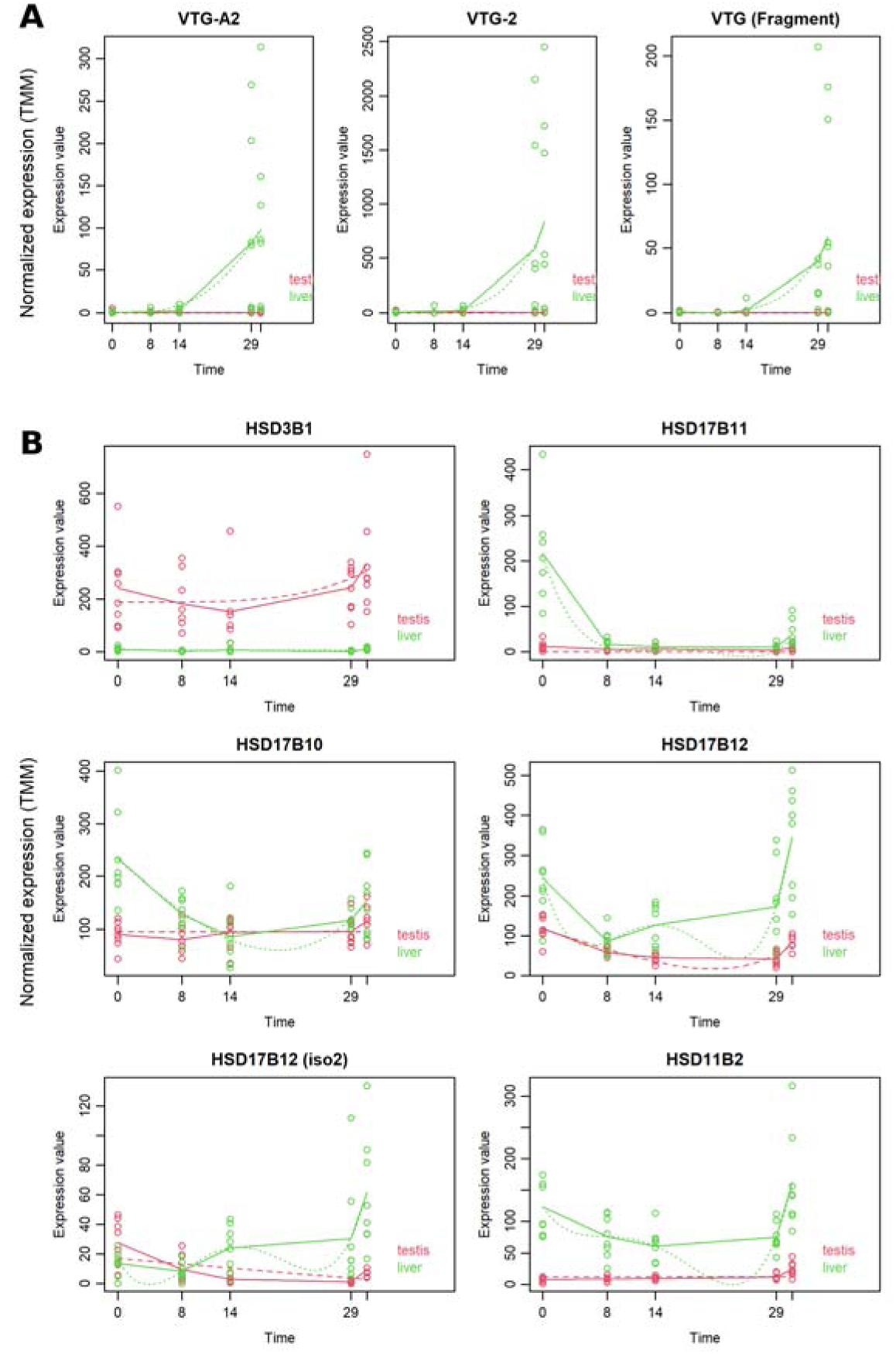
Vitellogenin and estrogen-modulating transcripts in male liver. (A) Liver (green) and testis (red) expression patterns for three vitellogenin transcripts: vitellogenin-A2, vitellogenin-2 and a vitellogenin fragment. (B) Expression patterns for steroid-modulating transcripts, including HSD3B1, HSD17B11, HSD17B10, HSD17B12, HSD17B12 isoform 2 and HSD11B2, in liver and testis. Points represent TMM-normalized expression values for individual biological replicates, with each biological replicate corresponding to one adult male snake; n = 8 animals per timepoint for each tissue. The experiment was performed once as a time-course RNA-seq experiment using independently sampled animals at each timepoint. Solid lines show observed mean expression for each tissue/timepoint, and dotted lines show fitted regression models. The x-axis is time in weeks from brumation onset; the post-arousal timepoint occurred at week 31 and is shown as the rightmost set of symbols in each panel. Transcripts were analyzed using time-course-aware negative-binomial regression in maSigPro on TMM-normalized counts with Benjamini–Hochberg correction for multiple testing; transcripts shown were significant at adjusted P < 0.05 in at least one analysis.

A coordinated estrogen-modulating panel accompanied this VTG response (Fig. 8B). 17β-Hydroxysteroid dehydrogenase 12 (HSD17B12), which converts the less reactive estrone to the more reactive estradiol (Luu-The et al., 2006; Blanchard, 2007), was upregulated as brumation progressed, while HSD17B11 and HSD17B10, which catalyze the reverse reaction (He et al., 2001; Brereton et al., 2001), were repressed. Aromatase transcripts were not detected in the liver, though very low-level expression cannot be excluded. 11β-Hydroxysteroid dehydrogenase 2 (HSD11B2), which inactivates glucocorticoids by converting cortisol to cortisone (Odermatt et al., 1999), was sustained-upregulated in liver but expressed only post-arousal in testis. 3β-Hydroxysteroid dehydrogenase / Δ –Δ -isomerase (HSD3B1), a hub enzyme in steroidogenesis (Simard et al., 2005), was high in testis throughout and absent from liver.

## Discussion

We propose a framework for understanding long-term ectotherm dormancy in which brumation is a dynamic physiological state regulated by two distinct transcriptional regimes: one temperature-associated and one starvation-associated. This framework predicts that different functional modules should be activated and deactivated at different points in the brumation cycle according to which regime is active, and that the two regimes should be distinguishable by their post-arousal recovery kinetics. Three high-level results are consistent with these predictions.

First, the overarching expression archetypes can be partitioned into two groups distinguishable by their post-arousal recovery kinetics, consistent with two separable regimes operating across the cycle. Second, liver shows a coordinated but selective transcriptional program in which pathways associated with lipid mobilization, alanine cycling and gluconeogenesis are maintained or upregulated within an overall suppressed metabolic background, consistent with a temperature-associated shift toward hepatic lipid mobilization that may reconcile the long-standing mismatch between the apparent stability of adipose stores and the energetic demands of the eight-month cycle. Third, testis shows sustained transcriptional suppression of meiosis, reproduction and DNA-metabolism gene sets that do not recover by the time the species enters its intense spring mating period, consistent with its dissociated reproductive pattern and passive sperm-storage strategy.

### Temperature- and starvation-associated transcriptional programs

The five overarching expression archetypes (Fig. 4) can be broadly interpreted along a thermal axis. Sustained downregulation and sustained upregulation, which persist post-arousal, are consistent with a starvation-associated response, since temperatures have already returned to active levels at the post-arousal sample but transcripts have not (Betts et al., 2002; Salem et al., 2007; Drew et al., 2008). The broad downregulation captured in combined cluster 1, spanning oxidative phosphorylation, beta-oxidation, and amino acid catabolism, likely reflects a passive, temperature-driven reduction in enzymatic demand rather than active transcriptional reprogramming and provides the metabolic backdrop against which the more selective, mid-brumation upregulations in hepatic lipid mobilization and gluconeogenesis stand out as coordinated, regulated responses. Downregulation with recovery and brumation-specific upregulation, which return toward pre-brumation values at arousal, track temperature. Because animals received no food from onset of brumation through post-arousal, starvation duration increased monotonically across all five timepoints even after temperatures had returned to active conditions. Thus, transcriptional programs that recovered following arousal despite continued starvation are more consistent with temperature-associated regulation, whereas programs that remained altered after warming are more consistent with prolonged energetic depletion. This separation is consistent with findings from a study in the closely related Checkered Garter Snake (*T. marcianus*) in which circulating metabolites and metabolic rate change with temperature during brumation (Holden et al., 2021) and further supported by our findings regarding the upregulated expression of a suite of heat shock proteins and antioxidant programs specifically during the cold period.

### A transcriptional program consistent with hepatic lipid mobilization at the coldest temperatures

A key result is the coordinated, mid-brumation peak in hepatic lipid mobilization (ATGL, FOXO1, PPARα, CPT1A, CPT2, ACSL4) coincident with the lowest chamber temperatures. This pattern provides a potential resolution to the apparent paradox in the existing physiological literature, in which adipose lipid stores in *T. s. parietalis* and other long-brumating squamates do not measurably decline across brumation but the energetics of the eight-month aphagous period clearly require some lipid utilization (Costanzo, 1985; Zani et al., 2012; Wilson, 2020). Hepatic lipid droplets, which are known to grow during feeding and shrink during fasting in *T. sirtalis* (Starck and Beese, 2002), therefore represent a plausible intracellular lipid source. Importantly, this pattern occurs alongside substantial downregulation of broader mitochondrial and metabolic gene sets, indicating that lipid mobilization represents a targeted metabolic priority rather than a global increase in metabolic activity. The gene-set enrichment analysis provides independent support for this interpretation. Liver cluster 2, which shows sustained downregulation, was enriched for broad fatty acid catabolic processes, organic acid catabolism, and oxidoreductase activity, which are the pathways expected to be upregulated if the animal were drawing on general adipocyte lipid reserves and processing a diverse pool of circulating fatty acids. The suppression of these pathways, against which the upregulation of hepatocyte-intrinsic droplet mobilization genes (ATGL, CPT1A, CPT2, ACSL4) stands out, argues that the liver is not responding to a systemic influx of adipose-derived fatty acids but is consistent with mobilization of a localized intracellular fuel reserve. The distinction is biologically meaningful: peroxisomal β-oxidation, enriched in the downregulated cluster, handles the very-long-chain and branched-chain fatty acids more characteristic of diverse adipose-derived substrates, whereas the gene panel evidence points to mitochondrial CPT1A/CPT2-mediated oxidation of hepatocyte-droplet fatty acids. The enrichment pattern therefore does not contradict the gene panel findings. Instead, it sharpens them, suggesting that broad systemic fat mobilization may be limited in favor of a small, targeted hepatic fuel source that could provide the minimal energy needed during the cold period. The relative maintenance of ApoB-100 expression alongside strong LDLR upregulation is consistent with continued hepatic VLDL output and potential increases in LDL recycling during the coldest period, suggesting that lipid mobilization includes both intracellular oxidation and lipoprotein-mediated redistribution. A similar pattern was reported in the Chinese soft-shelled turtle (*Pelodiscus sinensis*), where hepatocyte lipid droplets decreased in size during brumation and ATGL, PPARα, CPT1 and APOB-100 were transcriptionally upregulated (Huang et al., 2019). A multi-omic analysis of the Chinese alligator (*Alligator sinensis*) likewise showed hepatic upregulation of lipid catabolism without measurable change in adipose tissue size during brumation (Lin et al., 2020). Although hepatic lipid droplets were not measured directly here, the concordance between prior studies and the transcriptional patterns observed in *T. s. parietalis* are consistent with hepatic lipid-droplet mobilization as a testable mechanistic explanation.

These observations suggest a temperature-associated metabolic shift in which adipocyte mobilization may decrease below approximately 5 °C, while hepatic lipid droplets may become a more prominent intracellular lipid source. This interpretation is consistent with experimental data showing that brumating red-sided garter snakes held at 12 °C lose adipose mass while those at 4 °C do not (Wilson, 2020), and with the metabolite-level transition reported at ∼5 °C in *T. marcianus* (Holden et al., 2021). Such substrate partitioning would have a clear ecological logic: preserving adipose stores through brumation leaves them available to fuel the energetically costly month of post-arousal scramble-mating (Friesen et al., 2015) and the subsequent ∼17 km migration to summer feeding grounds (Aleksiuk and Stewart, 1971; Gregory and Stewart, 1975; Gregory, 2011).

The upregulation of ACAC and FASN and simultaneous repression of the lipid-droplet-storage enzymes ACAT2 and FITM2 is unusual. We interpret this not as evidence for net lipogenesis, but as a pattern compatible with diversion of newly synthesized fatty acids toward β-oxidation, VLDL export, or steroid synthesis rather than lipid-droplet storage. Experimental support for this interpretation would require lipid profiling and direct measurement of hepatocyte lipid-droplet dynamics during brumation in *T. s. parietalis* paralleling the histological work performed in *P. sinensis* (Huang et al., 2019).

### Transcriptional evidence consistent with alanine-supported gluconeogenesis

Despite earlier physiological work suggesting glycogen as a leading energy substrate during brumation in some squamates (Costanzo, 1985; Zani et al., 2012), no liver cluster in our data was enriched for upregulated carbohydrate-metabolism gene sets. The near-zero hepatic expression of HK1 throughout brumation, combined with the comparatively modest reduction in GLUT2, is more consistent with liver functioning as a glucose exporter rather than a glucose consumer during this period. This role would agree with the evidence for active gluconeogenesis via PCK1 and the Cahill-cycle program described below. The steeper decline in GBE1 relative to GAA similarly suggests a modest net shift toward glycogen catabolism over synthesis, though the overall carbohydrate signal is modest compared to the lipid-mobilization program. The mid-brumation upregulation of GDH1, GPT2L and SLC1A5 alongside PCK1, FOXO1, CREBBP, ACLY and MDH1 and the sustained-but-attenuated GLUT2 (Fig. 6) collectively provide transcriptional evidence consistent with hepatic operation of the Cahill cycle. Skeletal-muscle protein catabolism would supply alanine to liver for transamination to pyruvate, releasing ammonium for the urea cycle; pyruvate would then enter gluconeogenesis to be exported back to muscle as glucose. Importantly, this cycle is ATP-consuming in liver, and the parallel upregulation of CPT1A/CPT2 and ATGL provides a credible mitochondrial ATP source via fatty acid β-oxidation. The sustained upregulation of α-enolase (ENO1) is consistent with gluconeogenic activity given the near-absent HK1 expression, though its documented roles in thermal and hypoxic stress responses (Ji et al., 2016) make a dual function plausible and worth testing directly. Although protein catabolism is well established as a brumation-period energy source in long-brumating snakes (Aleksiuk and Stewart, 1971; Costanzo, 1985), our data add transcriptional evidence consistent with hepatic operation of the Cahill cycle, although biochemical validation in muscle and liver is needed before a mechanism can be assigned. As with lipid metabolism, these pathways appear to be selectively maintained rather than broadly upregulated, consistent with constrained metabolic function during prolonged cold exposure.

### Testis is reproductively quiescent but transcriptionally active

Testis showed sustained downregulation of clusters enriched for meiotic cell-cycle, reproduction and DNA-metabolism gene sets (Fig. 2; Table 1; Table S4). These transcripts dropped at brumation onset and did not recover by post-arousal. This is consistent with morphological and endocrinological work showing that *T. s. parietalis* testis is regressed during the spring mating period and contributes neither active spermatogenesis nor circulating androgens at that time (Crews et al., 1984; Krohmer et al., 1987). Together, our transcriptional data and the morphological literature reinforce that testis tissue is passively maintained across brumation with stored sperm produced during the previous summer/fall bearing the reproductive load through the spring (Aldridge, 1979; Clesson et al., 2002). The substantial brumation-specific upregulation of testis clusters enriched for translation, ribosome biology and mRNA processing (clusters 5, 12, and 13) suggests that, despite the absence of active spermatogenesis, testis maintains an active program of ribosome assembly, stored-mRNA processing and spliceosome-mediated splicing during brumation, presumably to preserve sperm viability over the long winter.

### Stress responses form a coordinated network

The integrated upregulation of HSPs, HIF1A, EP300, HYOU1, NRF2, the glutathione synthesis and recycling enzymes, SOD1 and PRDX1 indicates that *T. s. parietalis* mounts a multi-axis cellular stress response across brumation rather than a single cold-shock response (McCord and Fridovich, 1969; Lu 2009; Rhee et al., 2012). The post-arousal decline in most HSP transcripts is consistent with the reduction of long-term proteomic stress as temperatures rose, though expression remained elevated relative to most non-HSP transcripts at the post-arousal timepoint. Because post-arousal conditions in our study were more thermally stable than those during natural emergence conditions, free-ranging snakes encountering the variable thermal environment of spring may sustain higher HSP reliance than our data capture (Lutterschmidt et al., 2006). The further post-arousal increase in the ER-resident chaperone HYOU1 is consistent with elevated ER chaperone demand as translational activity resumed and may reflect a coordinated ER stress response during the metabolic transition out of brumation. The upregulation of HIF1A and EP300 during brumation is consistent with functional tissue hypoxia arising from the prolonged suppression of cardiac output and respiratory rate during dormancy, independent of environmental oxygen availability; HIF1A upregulation during normoxic torpor is documented in hibernating mammals and is thought to reflect reduced tissue oxygen delivery rather than ambient hypoxia (Morin and Storey, 2005; Maistrovski et al., 2012). The simultaneous downregulation of CAT and upregulation of GPX3/GLR1 is particularly notable. Two non-exclusive interpretations are consistent with our data: a temperature-associated switch in which catalase is replaced by the glutathione system below ∼10 °C, and a substrate-driven switch in which the rise in long-chain fatty acid utilization increases the demand for lipid-peroxidation–compatible antioxidants such as GPX3 (Yang et al., 2014, 2016; Ursini and Maiorino, 2020). Mixed reports of catalase activity at low temperature in other systems (Şahin and Gümüşlü, 2004; Chen et al., 2019) support the value of biochemical follow-up in *T. s. parietalis* tissue.

### Liver demonstrates brumation-specific upregulation of innate immune functions

Liver cluster 7, which shows brumation-specific upregulation (elevated expression during the cold period with post-arousal recovery), was enriched for 48 gene sets spanning innate and adaptive immune function, including complement activation, lymphocyte-mediated immunity, humoral immune response, B cell-mediated immunity, and inflammatory response. Notably, the top molecular function terms were endopeptidase inhibitor activity and peptidase inhibitor activity, enrichment profiles consistent with upregulation of serine protease inhibitors (serpins), which are predominantly synthesized in the liver and are central regulators of the complement cascade and acute-phase inflammatory response. Chromatin organization and chromatin remodeling were also significantly enriched, suggesting that the immune shift involves epigenetic reprogramming of immune gene expression rather than constitutive upregulation. The temperature-associated kinetics of cluster 7 indicate that this hepatic immune program is linked to cold exposure specifically, and not to the prolonged starvation that persists post-arousal.

One interpretation consistent with these patterns is that the liver selectively maintains or upregulates complement-based and humoral immune defenses during brumation as a temperature-tolerant alternative to cell-mediated immunity, which requires energy-intensive phagocytic and cytotoxic activity. Adaptive immunity in ectotherms is substantially impaired at low temperatures (Holden et al., 2021), while complement activity is generally less dependent on metabolically costly cellular processes. Serpin co-enrichment is coherent with complement upregulation: serpins such as C1-inhibitor and α2-macroglobulin modulate complement activation and limit excessive inflammation, a balance that would be important during prolonged dormancy when tissue damage must be minimized. Notably, this pattern contrasts with hibernating mammals, in which complement activity and hepatic C3 expression are reduced during torpor (Bouma *et al*., 2010), suggesting that the relationship between dormancy and hepatic immune function may differ fundamentally between endotherms and ectotherms.

### A vitellogenin response in male liver coincident with starvation

The most unexpected result is the strong upregulation of three vitellogenin transcripts in the male liver during late brumation, accompanied by a coordinated upregulation of the estradiol-favoring HSD17B12 and downregulation of the estradiol-degrading HSD17B11 and HSD17B10.

Aromatase was undetectable, suggesting that local androgen-to-estrogen conversion is unlikely to be the primary source of the estradiol signal; nonetheless, the steroid panel is consistent with conditions favoring estradiol presence in late-brumation liver. VTG is a yolk-precursor lipoglycophosphoprotein thought to be restricted to vitellogenic females and to males experimentally or environmentally exposed to estrogens (Wahli et al., 1981; Searle and Tata, 1981; Le Guellec et al., 1988; Lattier et al., 2001; Henry et al., 2009). Thus, VTG expression is not expected during natural physiology in adult males. Beyond its role as a yolk precursor, VTG has documented antimicrobial and antioxidant functions in invertebrates and fish (Shi et al., 2006; Seehuus et al., 2006; Havukainen et al., 2013), raising the possibility that its upregulation here reflects a stress-protective rather than strictly reproductive role, though this role remains speculative without additional experimental support.

Although the functional significance of this response is uncertain, we speculate that prolonged starvation may be linked to the long-standing observation that some male *T. s. parietalis* express female sexual-attractiveness pheromone at emergence (Mason and Crews, 1985; Shine et al., 2001; Parker and Mason, 2012). Both starvation status and expression of estrogen-modulating transcripts vary across the brumation cycle in ways that could plausibly intersect with pheromone biosynthesis. This possibility, though tantalizing, is limited in that this interpretation is based solely on sequencing data from male tissues and thus lacks protein-level support or direct estrogen quantification necessary to confidently establish a causal relationship. As this study sampled adult males only, it remains unknown whether a comparable vitellogenin response occurs in females during brumation, or whether the steroid-modulating pattern observed here is sex-specific. Together, these data identify an unexpected endocrine-like transcriptional response in male liver during late brumation that warrants targeted functional investigation.

### Testable predictions of the framework

The two-regime framework proposed here separates temperature-associated responses that recover after warming from starvation-associated responses that persist after arousal under continued aphagy. This framework generates specific, experimentally addressable predictions that distinguish it from a *post hoc* description of the data. First, if adipocyte mobilization is thermally gated near 5°C, lipid-droplet area in hepatocytes should decline across brumation at 4°C while adipocyte lipid content remains stable. Second, if hepatic lipid droplets are the primary intracellular lipid source during the cold period, circulating free fatty acid profiles should reflect a long-chain signature consistent with ATGL-driven triglyceride hydrolysis rather than adipose-derived fatty acid mobilization. Third, if the vitellogenin response is driven by prolonged negative energy balance rather than incidental estrogen exposure, VTG protein and circulating estradiol should covary with pre-brumation feeding state across individuals. Together, these predictions provide a roadmap for moving from transcriptional inference to mechanistic understanding of how long-brumating ectotherms partition energy across one of the longest periods of cold and aphagy documented in a squamate.

## Conclusions

The eight-month brumation of *T. s. parietalis* is, within the framework proposed here, a transcriptionally active state, not a state of bulk repression. Liver shows a coordinated transcriptional program consistent with temperature-associated hepatic lipid mobilization, potentially involving lipid droplets, alongside fatty acid β-oxidation and alanine-supported gluconeogenesis at the lowest brumation temperatures, accompanied by a coordinated stress response in which the antioxidant load shifts from catalase to glutathione. Testis shows sustained suppression of reproduction and meiosis through brumation, consistent with this species’ dissociated reproductive pattern, alongside upregulation of translation and RNA-processing modules likely required for stored-sperm maintenance. The unexpected late-brumation upregulation of vitellogenin and estradiol-favoring steroid-modulating enzymes in male liver opens a tractable hypothesis linking starvation status to the male sex-pheromone phenotype that has long been a puzzle of Thamnophis biology. The five-point temporal design should make this dataset a useful comparative reference for transcriptomic studies of other long-brumating squamates.

## Supporting information

Table S1: Experimental temperatures

Table S2: Sequence and mapping

Table S3: Liver enrichment results

Table S4: Testis enrichment results

Table S5: combined enrichment results

## Acknowledgements

All procedures were approved by the Oregon State University Institutional Animal Care and Use Committee under protocol 4818. Field research in Canada was conducted under the authority of Manitoba Wildlife Scientific Permit No. WB20333.

## Competing interests

The authors declare no competing or financial interests.

## Author contributions

D.L.H. conceived the study and wrote the original manuscript draft. D.L.H. and E.J.B. developed the methodology, performed the analyses, conducted the investigation. R.T.M. provided resources and supervision. All authors contributed to manuscript review and editing. Funding was acquired by R.T.M. and D.L.H.

## Funding

Financial support was provided in part by a National Science Foundation Graduate Research Fellowship to DLH (1840998).

## Data availability

Raw sequencing reads, processed count matrices and analysis scripts are deposited at NCBI SRA under BioProject PRJNA1473998 and on GitHub at https://github.com/hubertdl/Tsp_Brumation_Transcriptomics.

## Use of AI tools

During manuscript preparation and data analysis workflow development, the authors used ChatGPT (GPT-5.5, OpenAI) and Claude (claude-sonnet-4-6, Anthropic) to assist with code troubleshooting, language editing, and manuscript organization. All analyses, interpretations, and final text were reviewed and verified by the authors, who take full responsibility for the content of the manuscript.

## Supplementary Tables

Table S1: Brumation Conditions Table S2: Sequencing and Mapping

Table S3: Gene set enrichment for Liver Clusters Table S4: Gene set enrichment for Testis Clusters

Table S5: Gene set enrichment for combined tissue Clusters

---

**Fig. S1.**
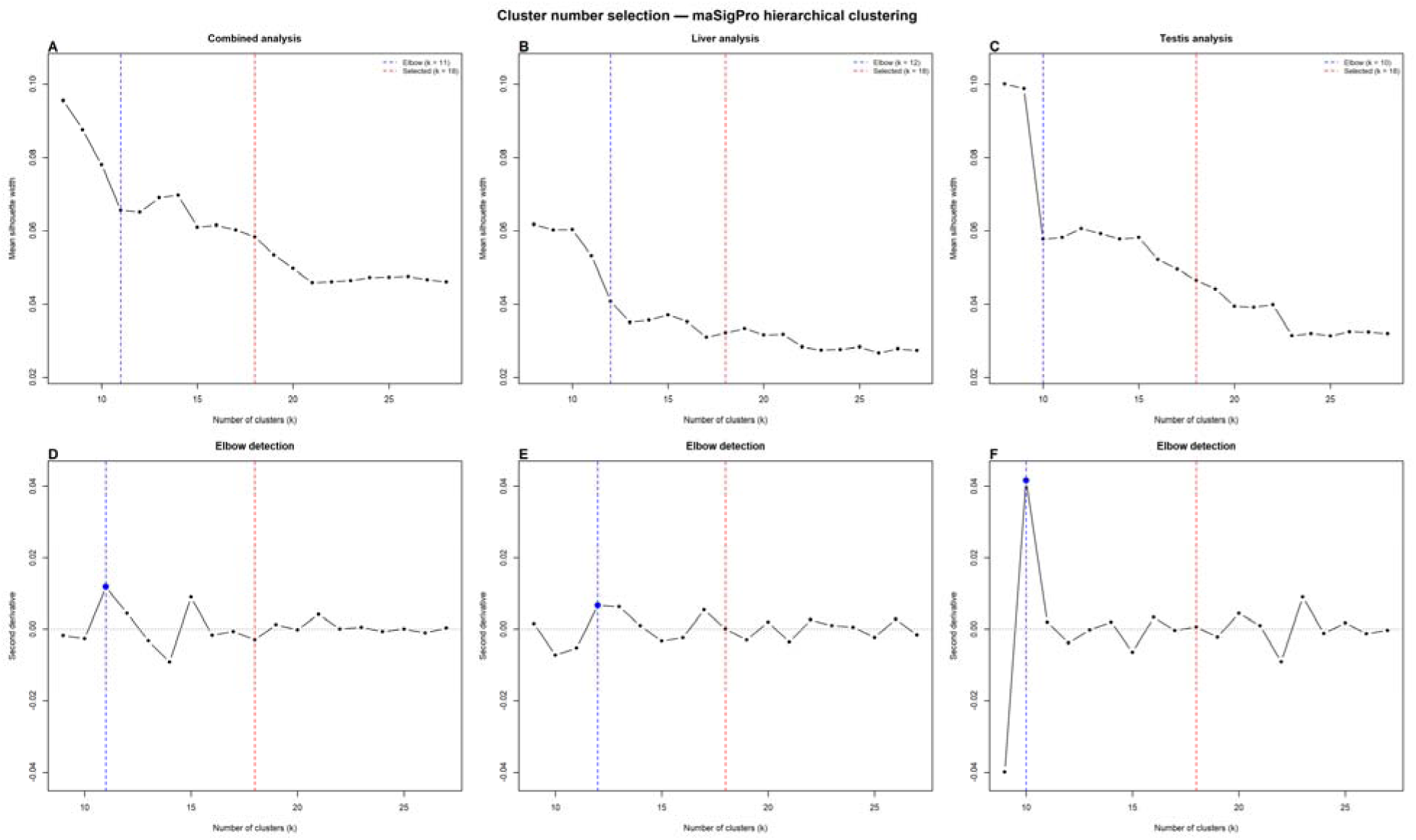
Cluster number selection for maSigPro hierarchical clustering. Mean silhouette width (A–C) and second derivative of silhouette scores (D–F) were evaluated across k = 8–28 clusters for the combined liver + testis (A,D), liver-only (B,E) and testis-only (C,F) maSigPro analyses. Analyses were based on TMM-normalized RNA-seq expression data from independently sampled adult male snakes, with n = 8 biological replicates per timepoint and five timepoints per tissue. Each biological replicate corresponds to one animal; no technical replicates are shown. The experiment was performed once as a time-course RNA-seq experiment. The elbow point, shown by the blue dashed line, identifies the initial inflection in the silhouette curve; k = 18, shown by the red dashed line, was selected as the minimum partition within the stable region of the curve that resolved biologically distinct expression sub-programs and reduced mean cluster size to fewer than 1,000 transcripts per cluster.

**Fig. S2.**
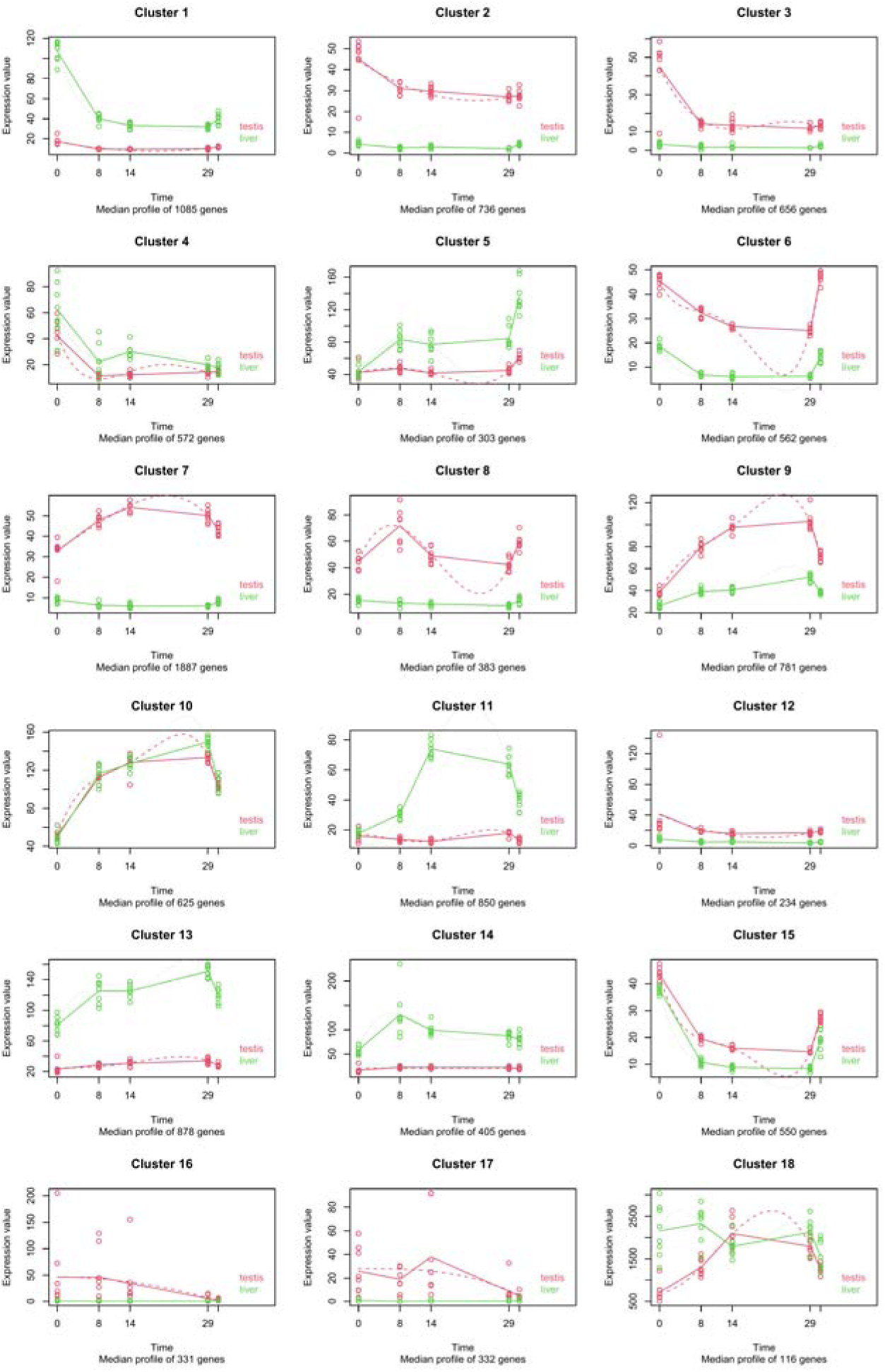
maSigPro time-course expression clusters in the combined liver + testis analysis. Hierarchical clustering of transcripts with significant temporal structure of 18 clusters in the combined liver + testis maSigPro analysis. Median-profile plots are shown for each cluster, with liver in green and testis in red. Points represent TMM-normalized expression values for individual biological replicates, with each biological replicate corresponding to one adult male snake; n = 8 animals per timepoint for each tissue. The experiment was performed once as a time-course RNA-seq experiment using independently sampled animals at each timepoint. Solid lines show observed median expression profiles for each tissue/timepoint, and dotted lines show fitted regression models. The x-axis is time in weeks from brumation onset; the post-arousal timepoint occurred at week 31 and is shown as the rightmost set of symbols in each panel. Clusters were identified using time-course-aware negative-binomial regression in maSigPro on TMM-normalized counts, with Benjamini–Hochberg correction for multiple testing; transcripts shown were significant at adjusted P < 0.05.

## Notes

### Competing Interest Statement

The authors have declared no competing interest.

